# PolyA-modulating antisense oligonucleotides reveal opposing functions for long non-coding RNA NEAT1 isoforms in neuroblastoma

**DOI:** 10.1101/2020.05.01.071696

**Authors:** Alina Naveed, Jack Cooper, Ruohan Li, Alysia Hubbard, Tao Liu, Steve Wilton, Sue Fletcher, Archa Fox

## Abstract

Many long non-coding RNAs (lncRNA) are highly dysregulated in cancer and are emerging as therapeutic targets. One example is NEAT1, which consists of two overlapping lncRNA isoforms, NEAT1_1 (3.7kb) and NEAT1_2 (23kb), that are functionally distinct. The longer NEAT1_2 is responsible for scaffolding gene-regulatory nuclear bodies termed paraspeckles, whereas NEAT1_1 is involved in paraspeckle-independent function. The NEAT1 isoform ratio is dependent on the efficient cleavage and polyadenylation of NEAT1_1 at the expense of NEAT1_2. Here we developed a targeted antisense oligonucleotide (ASO) approach to sterically block NEAT1_1 polyadenylation processing, achieving upregulation of NEAT1_2 and abundant paraspeckles. We have applied these ASOs to cells of the heterogeneous infant cancer, neuroblastoma, as we found higher NEAT1_1:NEAT1_2 ratio and lack of paraspeckles in high-risk neuroblastoma cells. These ASOs decrease NEAT1_1 levels, increase NEAT1_2/paraspeckles and concomitantly reduce cell viability in high-risk neuroblastoma specifically. In contrast, overexpression of NEAT1_1 has the opposite effect, increasing cell-proliferation. Transcriptomic analyses of high-risk neuroblastoma cells with altered NEAT1 ratios and increased paraspeckle abundance after ASO treatment showed an upregulation of differentiation pathways, as opposed to the usual aggressive neuroblastic phenotype. Thus, we have developed potential anti-cancer ASO drugs that can transiently increase growth-inhibiting NEAT1_2 RNA at the expense of growth-promoting NEAT1_1 RNA. These ASOs, unlike others that degrade lncRNAs, provide insights into the importance of altering lncRNA polyadenylation events to suppress tumorigenesis as a strategy to combat cancer.

## INTRODUCTION

Long non-coding RNAs (lncRNAs) have a key role in regulation of gene expression, both at transcriptional and post-transcriptional levels [1]. Dysregulation of lncRNA expression and function is well established as an important factor in the progression of cancer, developmental abnormalities and other diseases [2,3]. One such lncRNA is nuclear paraspeckle assembly transcript 1 (NEAT1), which occurs in two RNA isoforms expressed in the nucleus, NEAT1_1 and NEAT1_2 (3.7kb and 23kb, respectively, in human), both transcribed from a common promoter and overlapping at their 5’ end [4,5]. NEAT1 regulates gene expression in many ways, including transcription factor and miRNA sequestration, controlling pri-miRNA or pre-mRNA splicing, regulating RNA editing and enhancing protein stability [6]. Dysregulation of NEAT1 lncRNAs have previously been associated with various cancers [7], as well as neurodegenerative diseases such as amyotrophic lateral sclerosis (ALS) [8], Huntington’s disease [9] and Alzheimer’s disease [10].

NEAT1_2 has been extensively studied as an architectural lncRNA that is responsible for scaffolding subnuclear bodies called paraspeckles, that form when numerous RNA binding proteins bind to NEAT1_2 [11-13,4]. Paraspeckles are involved in the regulation of gene expression through subnuclear sequestration of component proteins and nuclear retention of RNAs [14]. It is only the production of NEAT1_2, not NEAT1_1, that provides a scaffold for protein binding and subsequent paraspeckle formation [12,13,15]. Paraspeckles have a distinct molecular organisation, with the 5’ and 3’ ends of NEAT1_2 at the periphery and the central sequences in the core of the 360nm-diameter bodies [16]. In contrast, NEAT1_1 is not sufficient for paraspeckle formation and has paraspeckle-independent roles [13,17]. Paraspeckle-independent foci containing NEAT1_1 have been dubbed microspeckles [17].

Dynamic changes in NEAT1 lncRNA levels are involved in tumorigenesis, where the ratio of NEAT1_1 to NEAT1_2 may be important. Various studies show NEAT1_1 primarily has a cell-proliferative role in cancer cells [18-21], whereas NEAT1_2 appearance can be associated with both tumour-suppressive and oncogenic activity [21-25]. In cancer, as well as other biological contexts, the overall transcription of NEAT1 is increased with stressors such as hypoxia, heat shock, mitochondrial stress and proteasome inhibition [26-28,18], driven by a number of upstream factors, including p53 [23,24]. The final ratio of the two NEAT1 isoforms relies on a series of protein binding events at the natural polyadenylation site (PAS) of the NEAT1_1 transcript [5,29].

NONO is an RNA binding protein found in paraspeckles, with high NEAT1_2-binding affinity [30]. A recent study demonstrates that NONO has a prognostic role in neuroblastoma, with higher levels of NONO correlating with high-risk neuroblastoma phenotype and poor patient outcome [31]. Neuroblastoma is the most common cancer seen in babies and infants and is a heterogenous disease with patients classified into low- or high-risk categories based on mutations and amplifications of a variety of different genes [32]. Of these, amplification of the oncogene MYCN is one of the clearest criteria for high-risk classification, with a 5-year event-free-survival rate of less than 50% in MYCN amplified patients [33,34]. MYCN maintains neuroblastoma cells in an undifferentiated state by dysregulating the ER pathway, amongst many other activities [35,36]. Consolidating its role as a master oncogenic regulator, overexpression of MYCN in neural crest cells leads to neuroblastic cell and neuro-ectodermal tumour formation [37]. Degree of differentiation is also a key factor in neuroblastoma, with high-risk neuroblastoma tumours containing a greater proportion of embryonic-type undifferentiated cells [34].

Existing neuroblastoma treatments can cause therapy-associated toxicity, and use of targeted therapies is an unmet clinical need [38]. In RNA therapeutics, the use of steric-blocking antisense oligonucleotides (ASOs) that modulate isoform expression of certain genes holds therapeutic potential in cancer, although most research is still at the preclinical stage [39]. In contrast, ASO-based RNA therapeutics in diseases such as spinal muscular atrophy and Duchenne muscular dystrophy have already been FDA-approved and are now being used to treat patients [40,41]. Here, we developed antisense oligonucleotides (ASOs) targeting the PAS of NEAT1_1, in an effort to promote greater transcriptional read-through to produce more NEAT1_2. We show that these ASOs lead to increased expression of NEAT1_2, decreased NEAT1_1 and also decreased cell growth of high-risk neuroblastoma cell lines. We observed that these ASOs lead to increased paraspeckle formation, and an upregulation of differentiation pathways in high-risk neuroblastoma cells with an associated reduction in neuroblastic phenotype, providing a proof of principle for the use of such ASOs in complex diseases.

## MATERIAL AND METHODS

### Patient Cohort Information

All patient RNA expression graphs and Kaplan-Meier Curves were obtained from R2: Genomics Analysis and Visualization Platform (Available http://r2.amc.nl; April 2020), with three neuroblastoma datasets. Datasets Versteeg (n=88) and Kocak (n=649) were selected because they had been previously examined for NONO and MYCN correlation by Liu and colleagues [31], and dataset SEQC (n=498) was chosen due to its large sample size and containing patient data from all 5 stages (1, 2, 3, 4, 4S). The Affymetrix Human Genome U133 Plus 2.0 Array was used in the Versteeg dataset, with probes 224566_at and 209757_s_at chosen to observe total NEAT1 and MYCN expression respectively of 88 patient tumour samples, with normalisation occurring using the MAS5.0 algorithm, recommended by the GCOS program of Affymetrix. The Agilent-020382 Human Custom Microarray 44k was used for both datasets Kocak and SEQC, where Kocak shows single-colour gene expression profiles from 649 neuroblastoma tumours. SEQC shows gene expression profiles obtained from 498 primary neuroblastoma samples. Probes UKv4_A_24_P566916 and UKv4_A_24_P94402 were chosen to study total NEAT1 and MYCN respectively for both datasets. In the Kocak dataset, 173 out of 649 patient survival data was unavailable, and hence not included in the Kaplan-Meier Curves (n = 476). All Kaplan-Meier Curves show overall survival probability of all patients, both relapse-free and otherwise. Two-sided Pearson’s coefficient was used to assess the correlation between expression levels of two genes. Kaplan-Meier curves were obtained using two-sided log-rank t-test.

### Cell culture and transfection preparation

Three neuroblastoma cell lines, KELLY, BE2C and SK-N-AS, were used in this study. KELLY and BE2C are cell lines that possess a genomic amplification of the MYCN gene, whereas the SK-N-AS cell line does not (Figure 1C). KELLY cells were grown in Gibco™ RPMI 1640 Medium, without phenol red (ThermoFisher, 11835055) with 10% Serana foetal bovine serum (FBS) (Fisher Biotec, FBS-AU-015) and 100U/mL penicillin-streptomycin (PenStrep) (ThermoFisher, 15140122), whereas the SK-N-AS, BE2C cell lines were grown in Gibco™ DMEM Medium high glucose, pyruvate media (ThermoFisher, 11995073) with 10% FBS and 100U/mL PenStrep. A549, HCT116 and U2OS cell lines were cultured in DMEM media with 10% FBS and 100U/mL PenStrep. All of these adherent cell lines were trypsinised for routine passaging using Gibco™ Tryple Express (ThermoFisher, 12604021). All cells were cultured in a 37°C incubator supplied with 5% CO_2_.

**Figure 1:**
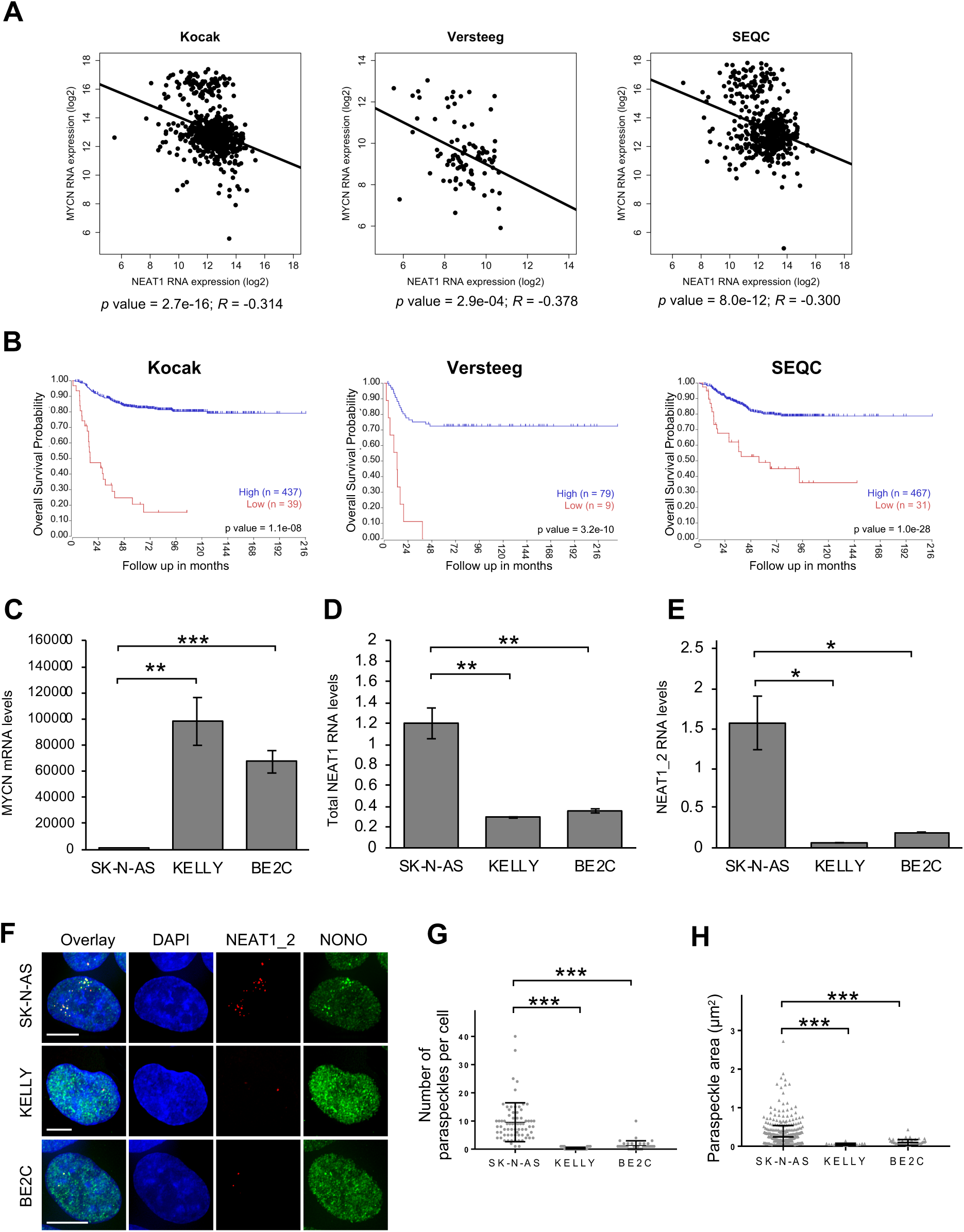
NEAT1 expression and paraspeckle abundance is lower in high-risk neuroblastoma patients and cell lines. (A) RNA expression levels of total NEAT1 and MYCN from the Versteeg, Kocak and SEQC neuroblastoma patient tumour gene expression datasets (https://hgserver1.amc.nl/cgi-bin/r2/main.cgi; Available April 2020). (B) Kaplan–Meier curves showing the probability of overall survival of patients according to total NEAT1 RNA expression in the same datasets as (A). (C-E) RT-qPCR data display relative (C) MYCN, (D) total NEAT1 and (E) NEAT1_2 RNA levels in the three neuroblastoma cell lines, SK-N-AS, KELLY and BE2C (n = 3, bars are SEM). (F) Fluorescence micrograph images of representative cells stained for NEAT1_2 and NONO in SK-N-AS, KELLY and BE2C neuroblastoma cell lines. DAPI (blue) stain cell nuclei, NEAT1_2 FISH (red) and NONO immunofluorescence (green). Scale Bar: 5µm. (G) Dot plot of the thresholded number of paraspeckles per cell and (H) paraspeckle area (µm^2^) for SK-N-AS, KELLY and BE2C neuroblastoma cell lines, stained for NEAT1_2 (e.g. As in F) (n ≥ 40 cells per cell line). (*P<0.05, ** P<0.001, ***P<0.0001)

All transfections were performed in a forward transfection manner, with cells plated in RPMI (4% FBS) or DMEM (4% FBS) without PenStrep one day prior to transfection. 96-well plates were used in cell viability and cell confluence assays, cells were plated in 100 μL volumes per well at a cell density of 5×10^3^ cells per well for KELLY, 1.5×10^3^ per well for BE2C and 4 ×10^3^ cells per well for SK-N-AS cells. 12-well plates were used to obtain RNA, protein or microscopy samples, cells were plated in 1 mL volumes per well at a cell density of 1×10^4^ cells per well for KELLY, 5×10^3^ per well for BE2C and 1×10^4^ cells per well for SK-N-AS cells.

### Antisense Oligonucleotide and Plasmid transfections

ASOs with two different chemically modified backbone moieties were used in this study; 2’O methyl modified bases on a phosphorothioate backbone (2’OMe) and phosphorodiamidate morpholino oligomers (PMO). In both cases the ASOs were fully modified along their entire length so as not to stimulate RNaseH mediated degradation of the target RNA, as is the case with gapmers, but rather to inhibit the cleavage and polyadenylation proteins binding to the target [42]. All 2’OMe ASOs were synthesised in-house on an Akta-OligoPilot plus 10 using the 1 micromol synthesis protocol, and stored at -20°C. The equivalent sequences were synthesized as PMOs and were supplied from Genetools LLC (Philomath, Oregon, USA). The sense strand DNA phosphodiester leash sequences were supplied by Integrated DNA Technologies, Inc (Coraville, Iowa, USA). To ensure efficient annealing of the PMO with its DNA leash, all morpholino solutions were incubated to 95°C for 2 mins, with 5°C temperature decrements for 2 mins each occurring thereafter until 20°C was reached, prior to every transfection. Lipofectamine ® 3000 (ThermoFisher, L3000015) and Lipofectamine ® RNAiMAX (ThermoFisher, 13778150) were the main transfection reagents used in this study. All transfection mixtures were made up in serum-reduced Gibco™ Optimem (ThermoFisher, 11058021), with 10 µL volumes used for transfection into 96-well and 100 µL volumes used for transfection into 12-well plates. All ASOs used for transfections were taken from a 10 µM stock, with transfections conducted using Lipofectamine ® RNAiMAX for both 2’OMe ASOs or PMOs. Final concentrations of oligonucleotides in 96-well or 12-wells varied between 25 nM to 100 nM, as indicated in results and figure legends. For all sno-control or sno-NEAT1_1 plasmid-only transfections, Lipofectamine ® 3000 was used, with 1 µg transfected into cells in 12-well plates, as per the manufacturer’s instructions. For dual oligo and plasmid transfections conducted in 12-well plates, Lipofectamine ® RNAiMAX only was used to transfect both 1 µg of plasmid and PMOs in the same transfection mix, with a final PMO concentration of 50 nM.

### RNA Isolation and RTqPCR

Cells were lysed with Nucleozol™ (Macherey-Nagel, 740404) and RNA extraction was conducted according to the manufacturer’s instructions. For the semi-extractible step (Figure 5D), samples were placed in Eppendorf Thermomixer® C for 1,000 rpm at 65°C for 10 minutes after RNase-free water was added to Nucleozol-lysed samples. To visualise pellets more easily in subsequent ethanol washes, GlycoBlue™ coprecipitant (ThermoFisher, AM9516) was added during the isopropanol addition steps. RNA samples were reverse transcribed using the QuantiTect Reverse Transcription Kit™ (Qiagen, 205314). qPCR was conducted using custom-made primers (Integrated DNA Technologies, Inc, Coraville, Iowa, USA) (see Table S1 for primer sequences), and SensiMix SYBR No-ROX Kit™ (Bioline, QT650-20), using RPLP0 as a reference gene. qPCR reactions were conducted in Qiagen Rotogene Q, with Ct values acquired using a threshold of 0.075 for all sample sets and 2^-ΔΔCt^ method used for relative quantitation. The comparative quantitation feature of the Rotogene Q software was used to obtain amplification and take-off values used to determine ratios of NEAT1_1 and NEAT1_2.

**Figure 2:**
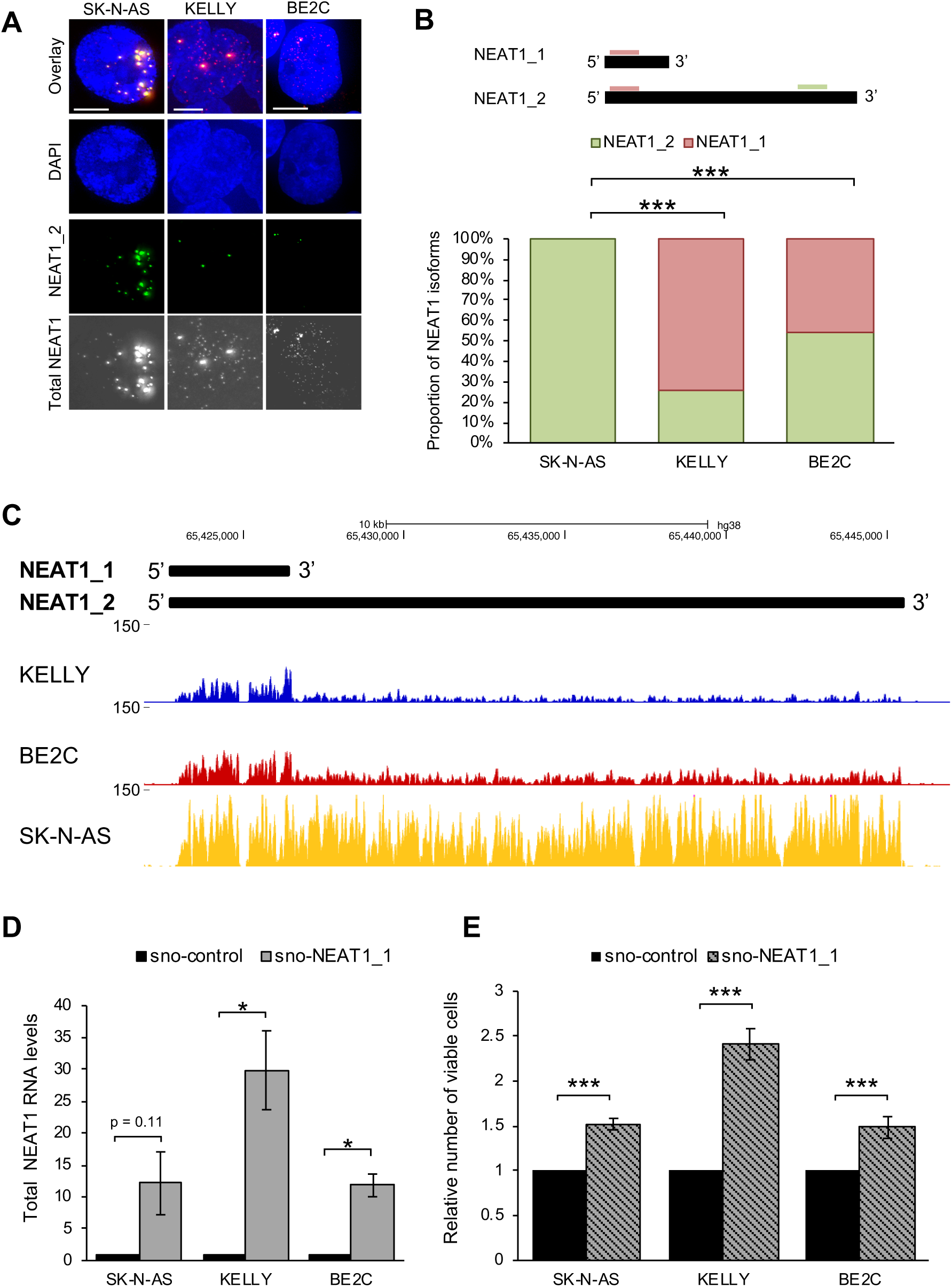
NEAT1_1 is expressed in high-risk neuroblastoma cell lines, and can drive proliferation in neuroblastoma cells. (A) Fluorescence micrograph images of representative cells stained for NEAT1_2 (green) and total NEAT1 (greyscale) in SK-N-AS, KELLY and BE2C cell lines. DAPI (blue) stain cell nuclei, NEAT1_2 FISH (red) and total NEAT1 FISH (green). Scale Bar: 5µm. (B) Schematic of NEAT1 isoforms showing location of total NEAT1 (pink) and NEAT1_2 (green) primers used for RT-qPCR to comparatively quantitate NEAT1_1 and NEAT1_2 RNA expression levels in SK-N-AS, KELLY and BE2C cell lines (SEM for KELLY = 2.48% and for BE2C = 3.35%). (C) RNA-seq coverage across the NEAT1 gene in KELLY, BE2C and SK-N-AS cell lines. Scale bar is 150 reads/bp for all cell lines. (D) RT-qPCR data indicating relative total NEAT1 levels in SK-N-AS, KELLY and BE2C cells 48h post-transfection with sno-control, or sno-NEAT1_1 plasmids, as indicated (n = 3). (E) Cell viability was measured using the Cell Titer Glo assay 48h post-transfection of SK-N-AS, KELLY and BE2C cells with sno-control plasmid or sno-NEAT1_1 plasmid (n = 3). (*P<0.05, **P<0.001, ***P<0.0001).

**Figure 3:**
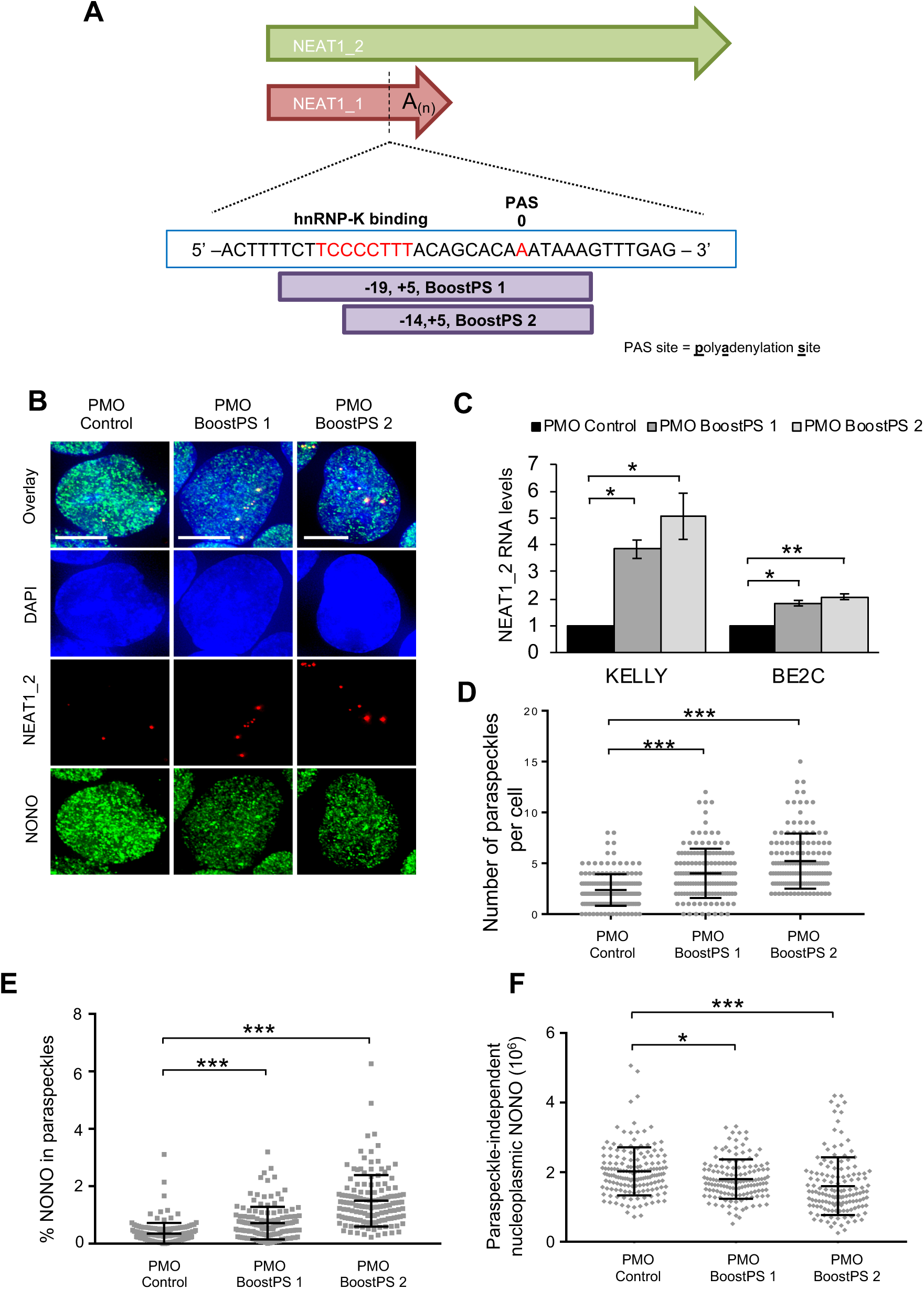
Morpholinos (PMOs) targeted to the polyadenylation site within NEAT1_1 lncRNA can increase NEAT1_2 levels, resulting in increased paraspeckle formation. (A) Schematic showing the binding sites of BoostPS 1 and BoostPS 2 PMOs at the polyadenylation site (PAS) sequence within the NEAT1 RNA transcript. (B) Fluorescence micrograph images of representative cells stained for NEAT1_2 and NONO in KELLY cells transfected with 25nM PMO: Control PMO, left, Boost PS 1 PMO (middle) and Boost PS 2 PMO (right). DAPI (blue) stain cell nuclei, NEAT1_2 FISH (red) and NONO immunofluorescence (green). Scale Bar: 5µm. (C) RT-qPCR data indicating relative NEAT1_2 levels in KELLY and BE2C cells treated with 25 nM Control or BoostPS PMOs for 48h (n = 3). (D) Dot plot of the thresholded number of paraspeckles per cell for KELLY cells transfected with 25nM Control or BoostPS PMOs and stained for NEAT1_2 (eg. As in B). (E-F) Percentage of nuclear NONO fluorescence detected within paraspeckle foci (E), or out with paraspeckle foci (F), as defined by red NEAT1_2 thresholded regions, in KELLY cells transfected with 25nM Control or BoostPS PMOs (n ≥ 140 cells). (*P<0.05, **P<0.001, ***P<0.0001)

**Figure 4:**
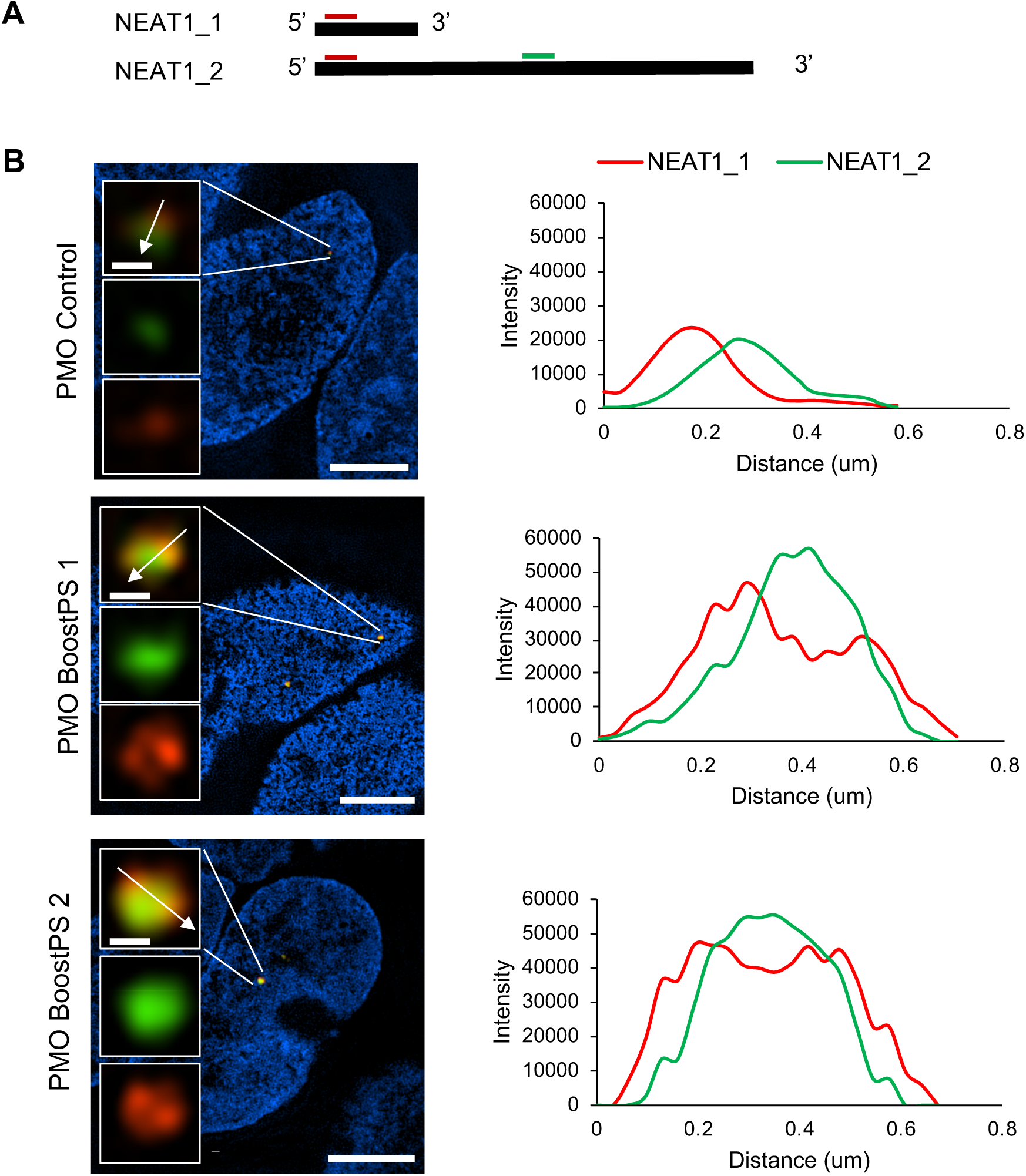
Increased NEAT1_2 RNA expression with BoostPS PMO treatment in KELLY cells leads to formation of traditional core-shell paraspeckles. (A) Schematic showing the approximate regions of NEAT1_1 and NEAT1_2 targeted by the dual colour FISH probes used in (B). (B) Structured illumination microscopy was conducted in KELLY cells transfected with 50 nM Control or BoostPS PMOs and stained with dual-colour FISH probes against NEAT1_1/NEAT1_2 5’ end (red) and NEAT1_2 (green) and DAPI (blue). Images on the left show representative fluorescence micrograph images of a single section captured in the middle of a paraspeckle, with intensity profiles of the white arrow lines depicted on the right. Scale Bar for cell nuclei: 5 µm; Scale bar for paraspeckle inset images: 0.5 µm.

**Figure 5:**
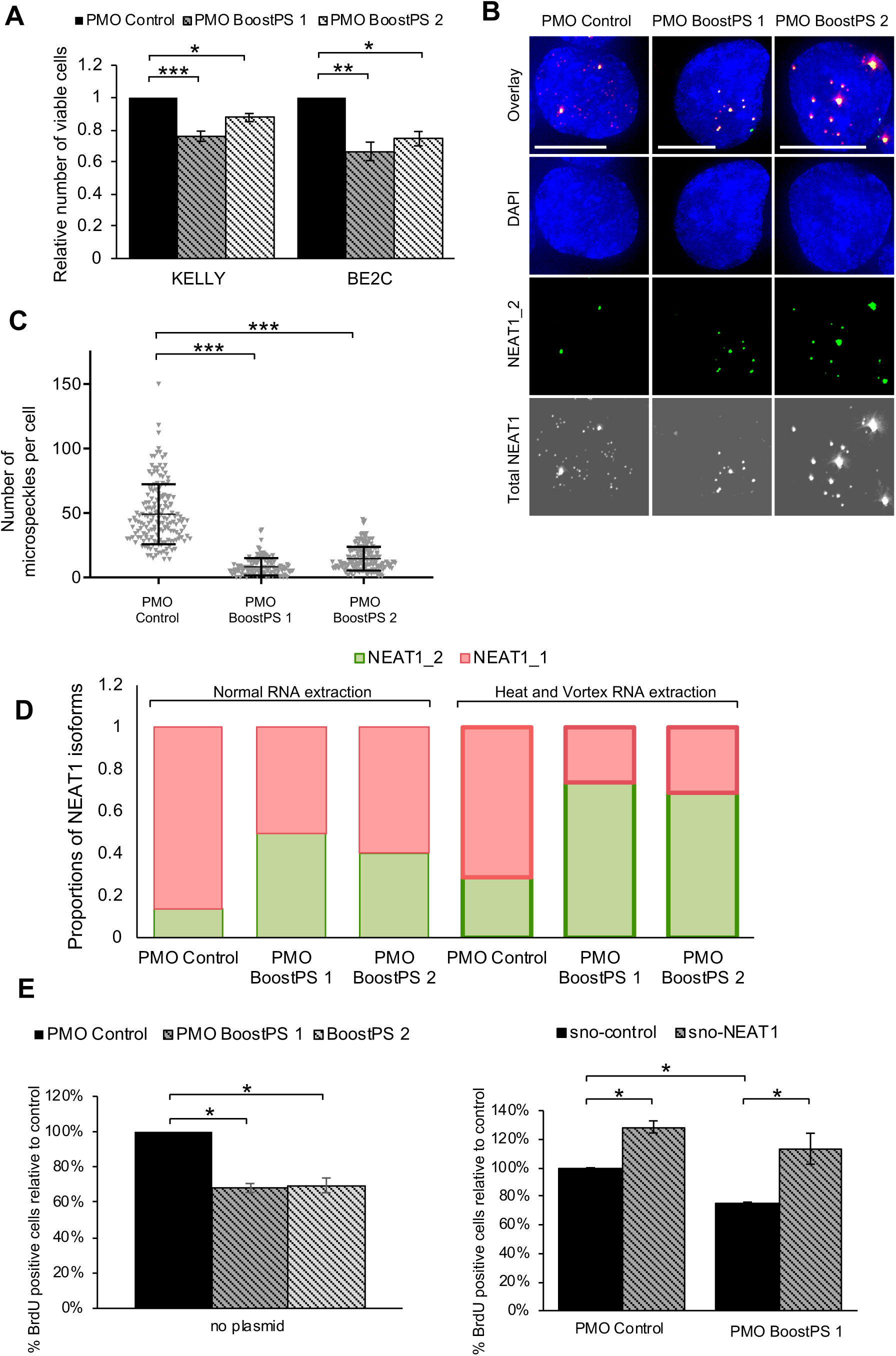
High-risk neuroblastoma cells treated with BoostPS PMOs show decreased NEAT1_1 RNA expression levels, along with reduced cell viability. (A) Cell viability was measured using the Cell Titer Glo assay, 6 days post-transfection of KELLY (left) or BE2C cells (right) with 25 nM Control or BoostPS PMOs (n = 3). (B) Fluorescence micrograph images of representative cells stained for total NEAT1 (greyscale) and NEAT1_2 (green) RNA using FISH in KELLY cells transfected with control (left), Boost PS 1 (middle) or Boost PS 2 PMOs (right). DAPI (blue) stain indicates cell nuclei. Scale Bar: 5µm. (C) Dot plot for the thresholded number of microspeckles per cell in KELLY cells transfected with 25 nM Control or BoostPS PMOs (treated as in (C)). (D) RT-qPCR data indicating proportions of NEAT1_1 and NEAT1_2, calculated using comparative quantitation, for KELLY cells 48h post-transfection with 50 nM Control or BoostPS PMOs in samples extracted using normal or heat-and-vortex RNA extraction protocol (n = 2). (E) Percentage of KELLY cells incorporating BrdU in a 2h pulse-label, 4 days post-transfection of 50 nM Control or BoostPS PMOs. Graph on the left indicates cells with PMO transfection only. Graph on the right shows data for cells co-transfected with either Control or BoostPS 1 PMOs with sno-control or sno-NEAT1_1 plasmids (n = 3). (*P<0.05, **P<0.001, ***P<0.0001)

### Immunofluorescence and RNA fluorescence in situ hybridization

Cells were grown on and fixed onto coverslips (diameter 18mm, thickness 1.5; Schott, G405-15) using 4% paraformaldehyde in PBS. Immunofluorescence involved permeabilisation in 1% Triton X-100 diluted in PBS for 10 minutes, followed by rinsing three times in PBS. Epitope detection was conducted with a monoclonal mouse antibody against NONO [43] at 1:500 dilution in PBST, followed by three PBS washes, then secondary FITC anti-mouse antibody (Jackson Laboratories, 115-095-072), at 1:250 dilution in PBST. Coverslips were then hybridised overnight with fluorescence in situ hybridisation probes (FISH), according to the manufacturer’s (i.e. Stellaris) instructions.

The probes used in this study include Human NEAT1 5’ Segment with Quasar ® 570 Dye (Stellaris, VSMF-2246-5), Human NEAT1 Middle Segment with Quasar ® 570 Dye (Stellaris, SMF-2037-1) and Human NEAT1 Middle Segment with FAM Dye (Stellaris, VSMF-2248-5) probe sets. For FISH without prior immunofluorescence, permeabilisation was conducted by incubating coverslips in 70% ethanol overnight at 4°C. For dual-probe FISH experiments to detect both total NEAT1 and NEAT1_2 RNA, Stellaris® FISH Probes Human NEAT1 5’ Segment with Quasar ® 570 Dye and Human NEAT1 Middle Segment With FAM Dye respectively were used, with both probes added in the same hybridisation buffer reaction.

### Cell viability assays

Transfections were conducted one day after cells were plated, with fresh media supplied (50 µL of 15% FBS RPMI or DMEM media per well) every second day following transfection. For Cell Titer Glo assays, cells were plated into white-coloured opaque 96-well plates (with at least 3 and maximum 6 technical replicates used per biological replicate). Six days post transfection, cells were incubated with 10 µL Cell Titer Glo (Promega®) per well and plates were shaken for 2 minutes. Luminescence activity was measured and recorded using the Fluostar Optima Instrument.

### Cell confluence assays

For cell confluence experiments, cells were plated into clear nuncleon-coated 96-well plates (with 2-3 technical replicates used per biological replicate) on Day 0 and left in the incubator for 24 hours. Immediately after transfection, plates were then placed in an Incucyte™ S3 Live-Cell Imaging System and each well was scanned using the Phase channel at 4x objective, once daily for 6 days. All Incucyte data analyses for KELLY cells were conducted using default settings, with Phase Segmentation Adjustment set at 0.5.

### BrdU cell replication assay

Cells were plated onto coverslips in 12-well plates. The following day, cells were transfected with either PMO only or PMO and plasmid and cultured for a further 4 days. 2 hours prior to fixation, media was replaced with media containing 10 nM 5-Bromo-2′-deoxyuridine (BrdU) (Merck, B5002-100MG). Coverslips were then fixed, and cells were permeabilised in 1% Triton X-100/PBS for 10 minutes at room temperature. Coverslips were rinsed twice in 500 uL of PBS, swirling coverslips in PBS for 10 seconds each time. The coverslips were incubated in 1 M HCl for 30 minutes, rinsed twice in PBS and then incubated in 0.1 M sodium borate for 30 minutes. After blocking in 5% goat serum in PBST (50 µL per coverslip), immunofluorescent staining was carried out using Anti-BrdU antibody (Abcam, ab8955) at 1:500 dilution in PBST, followed by anti-mouse Cy™5-conjugated secondary antibody (Jackson Laboratories, 115-175-072) at 1:300 dilution in PBST. Cells were imaged on the Deltavision microscope using 60X objective (see below). Cells that had incorporated BrdU into their DNA during replication (labelled by Cy5) were measured and expressed as a value relative to the total number of nuclei, as measured by DAPI staining.

### Microscopy and image analysis for paraspeckle data

All images were acquired on a Deltavision Elite microscope (GE) using a 60X or 100X objective. For subsequent paraspeckle counting, the Nikon NiS Elements software (Version 4.3, Nikon, Tokyo, Japan) was used, with ensured size/area calibration implemented when viewing and analysing Deltavision acquired images with NiS. Acquisition parameters were kept consistent for samples compared between each other. All intensity thresholds (SumGreen for NONO and SumRed for NEAT1_2 intensities) were set the same across all groups during paraspeckle number, paraspeckle area, nucleoplasmic NONO and paraspeckle-NONO localisation quantitation in Nikon NiS Elements software. Unbiased raw data were obtained and analysed, with dot plots generated in GraphPad Prism 7.

### Structured Illumination Microscopy (SIM) and image processing

Images were acquired using a Nikon N-SIM microscope (Nikon, Tokyo, Japan) using a 100x objective (SR Apo TIRF objective, 1.49NA, automatic correction collar) and Andor iXON Ultra EMCCD camera, in 3D stack SIM mode. Stack volumes were determined visually, with top and bottom slices defined by reduction in corresponding paraspeckle fluorescence. Z step size was set to 120nm. Images were reconstructed using the SIM algorithm in NIS Elements (Version 5.2, Nikon, Tokyo, Japan). Intensity profile graphs obtained using NIS Elements (Version 4.3, Nikon, Tokyo, Japan).

### Protein lysis and Western Blotting

Cells in 12-well plates were lysed with 150 µL of 2x Protein lysis buffer (160 mM Tris pH 6.8, 4% SDS, 15% glycerol, 0.0006% bromophenol blue, and 0.004% β-mercaptoethanol), and protein lysates were heated to 95°C for 10-15 min. Lysates were loaded onto Mini-PROTEAN ® TGX™ Precast Protein Gels (Bio-Rad, 4561086) with Precision Plus Protein™ All Blue Prestained Protein Standards (Bio-Rad, 1610373) loaded alongside. Gels were run in Tris/Glycine Buffer (Bio-Rad, 1610771) at 200V for 30-40 min. Transfer was conducted with Trans-Blot ® Turbo™ Mini Nitrocellulose Transfer packs (Bio-Rad, 1704158) using the Trans-Blot ® Turbo™ Transfer System. Primary antibodies used were N-Myc Antibody [B8.4.B] (Santa Cruz, sc-53993), ALK [D5F3] (Cell Signalling Technology, #3633) and Anti-PHOX2B antibody [EPR14423] (Abcam, ab183741). All primary antibodies were diluted 1:500 in 5% milk PBST. The secondary (horseradish-peroxidase conjugated) antibodies used were Goat Anti-Rabbit IgG H&L (Abcam, ab97051) and Goat Anti-Mouse IgG H&L (HRP) (Abcam, ab97051), and were diluted 1:10,000 in 5% milk PBST. Luminata Crescendo Western HRP substrate (Merck, WBLUR0100) with image capture by Bio-Rad Chemidoc was used to visualise protein bands. Bio-Rad Imagelab Version 5.2 was used to quantify total protein levels and the intensity of the chemiluminescent bands. For normalisation, Imagelab software was used to determine the amount of total protein in each lane, and intensity of chemiluminescent bands were normalised back to this value. To determine chemiluminescent band sizes, the sizes of the bands of the BioRad All Blue Prestained Protein Standards ladder were visualised on the membrane images acquired on the Chemidoc, and chemiluminescent bands sizes were determined in relation to the standard ladder bands.

### KELLY PMO transfections and RNA-sequencing

KELLY cells were plated at a density of 3×10^5^ cells per well in 6-well plates, in 2 mL 4% FBS-RPMI media per well, and placed in a 37°C incubator overnight. After annealing the PMO:leash lipoplexes, transfections of the PMOs at a concentration of 50 nM was conducted using Lipofectamine ® RNAiMAX and cells were left in transfection media for 48 hours. Media was removed, cells were washed with 2 mL PBS twice, and lysed for RNA lysates, at outlined above. RNA isolations were conducted including a semi-extractability step, as described above. Samples were sent to the Australian Genome Research Facility (AGRF), and RNA quality was confirmed using a Bioanalyser (Perkin Elmar, MA, USA) prior to RNAseq. Whole transcriptome library preparation using the TruSeq Stranded Total RNA Library Prep Kit (Illumina, CA, USA) and ribosomal RNA depletion with the Ribo-Zero-Gold kit (Illumina, CA, USA) was performed. Sequencing was performed using an Illumina HiSeq 2500 (Illumina, CA, USA) to generate 100 bp single end reads, resulting in an average 25 million reads per sample. For RNA-sequencing of the three neuroblastoma cell lines, a similar approach was used, with the following changes: no transfections, no semi-extractability step, and 50 bp single end reads.

### Bioinformatics

Raw sequencing files were quality checked using FastQC (0.11.7), with all files passing. No adapter contamination was identified, and reads were not trimmed. Transcript quantification was performed with salmon (0.8.2), using an index constructed from the Ensembl GRCh38.93 annotation. Transcript abundance was summarised to gene-level counts and imported into R using tximport (1.4.0). Differential expression analysis was performed using DESeq2 (1.16.1) and the default parameters (alpha = 0.1). Three primary comparisons were performed: control (4) vs BoostPS1 (4), control vs BoostPS2 (4) and control (4) vs combined BoostPS1 and BoostPS2 (8). The heatmap of differentially expressed genes includes all genes that showed a significant change (p_adj_ < 0.1) and was constructed using the heatmap.2 function from R package gplots (3.0.1). Gene ontology analysis was performed using GSEA (3.0) with 1000 gene set permutations and the msigdb ‘BP: GO biological process’ gene set. The gene ontology network was constructed using the Enrichment Map plugin for Cytoscape (3.5.1), with cut-offs: p < 0.05, FDR < 0.1 and edge > 0.375. RNA-seq coverage tracks were generated using HISAT2 (2.1.0) for alignment to a GRCh38 genome index, samtools (1.4.1) for conversion to bam files and bamCoverage (3.1.2) for conversion to bigwig and upload to the UCSC genome browser (https://genome.ucsc.edu/index.html).

### Statistical Analyses

All graphs show experiments from three biological replicates (n = 3), unless otherwise stated. All dot plot datasets were generated and analysed using Graph Prism 7. A two-tailed unpaired Student t-test was conducted assuming unequal variance for all dot plots. All dot plots with error bars show mean with standard deviation, unless otherwise stated. All column graphs were generated and analysed using Microsoft Excel 2013. A two-tailed unpaired Student t-test was conducted assuming equal variance for all RTqPCR datasets. For cell viability data, cell confluence data and BrdU data, a two-tailed unpaired Student t-test was conducted assuming unequal variance. For all Western Blot datasets, a two-tailed unpaired Student t-test was conducted assuming equal variance. All column graphs with error bars show mean with standard error mean, unless otherwise stated.

## RESULTS

### NEAT1 expression levels and paraspeckle abundance is lower in high-risk neuroblastoma

We first examined NEAT1 expression levels in three different neuroblastoma patient cohorts, and observed an inverse correlation between MYCN and NEAT1 levels, suggesting high-risk patients express lower total NEAT1 (Figure 1A). Kaplan-Meier curves showed that patients with poor outcomes express lower levels of NEAT1 (Figure 1B). Next, we examined NEAT1 expression in three neuroblastoma cell lines; non-MYCN amplified low-risk SK-N-AS and two MYCN amplified high-risk KELLY and BE2C cells (Figure 1C). Given both NEAT1 isoforms precisely overlap, we conducted RT-qPCR using primers amplifying ‘total’ NEAT1 or only NEAT1_2 (schematic as per Figure 2B). The MYCN amplified KELLY and BE2C cells exhibit lower RNA levels of total NEAT1 (Figure 1D) and NEAT1_2 (Figure 1E) compared to non-MYCN amplified SK-N-AS cells, mirroring the pattern seen in neuroblastoma patient tumour samples. These data propose a possible inverse correlation between NEAT1 and MYCN, with commensurate risk status, in neuroblastoma.

We next visualised paraspeckles using fluorescence in situ hybridisation (FISH) against NEAT1_2, in conjunction with immunofluorescence against NONO. SK-N-AS cells, with high NEAT1 levels, have large and prominent paraspeckles as marked by NEAT1_2 and NONO co-localisation, whereas KELLY and BE2C had smaller and fewer paraspeckles (Figure 1F). Quantitation of the images confirmed this, with SK-N-AS paraspeckle number (mean ± SD; 9.54 ± 0.82) greater than KELLY (0.25 ± 0.05) and BE2C (1.28 ± 0.27) (Figure 1G). The average paraspeckle area was also larger in SK-N-AS cells (Figure 1H). These data support NEAT1_2 and paraspeckles being associated with potential tumour-suppressive function, as shown previously in multiple studies [21,22].

### Microspeckles in high-risk neuroblastoma cells have a cell-proliferative role

Next, NEAT1_1 within the nuclei of the neuroblastoma cells was visualised. Previously, we reported paraspeckle-independent ‘microspeckles’ of NEAT1_1 foci occurring in osteosarcoma cells [17]. To examine microspeckles in neuroblastoma cells, we used two sets of FISH probes: a set with red fluorophores targeted to the NEAT1 5’-end (detecting both NEAT1_1 and NEAT1_2) and a set with green fluorophores targeted only to NEAT1_2 (schematic as per Figure 4A). Figure 2A shows KELLY and BE2C exhibit numerous microspeckles (greyscale in lower panel and red in overlay image), which are not coincident with the rare, green paraspeckles. Remarkably, SK-N-AS cells show apparent complete overlap of NEAT1_2 (green) and NEAT1_1 (red) foci (Figure 2A, overlay image), suggesting SK-N-AS has very low, or undetectable NEAT1_1. RT-qPCR confirmed these findings; SK-N-AS cells have higher NEAT1 levels overall (Figure 1D/E), yet lack NEAT1_1 (Figure 2B), whereas KELLY and BE2C cell lines express lower NEAT1 levels overall (Figure 1D/E) however the NEAT1 they do express consists largely of NEAT1_1 (Figure 2B). These findings were confirmed with RNA-seq: KELLY and BE2C have more reads mapping across NEAT1_1 and lower mapping across NEAT1_2, whilst SK-N-AS demonstrates high read density across the entirety of NEAT1_2, indicating an absence of NEAT1_1 (Figure 2C).

The high levels of NEAT1_1 in the more aggressive, high-risk neuroblastoma cells suggests NEAT1_1 has an oncogenic role here, as has been described in other cancers in other studies [21,19]. To test this, we overexpressed NEAT1_1, using a ‘sno’-vector designed to ensure nuclear localisation of the RNA [44] (Figure S1A). We transfected sno-control or sno-NEAT1_1 plasmid in all three neuroblastoma cell lines, and observed an increase in total NEAT1 RNA expression levels (Figure 2D). FISH to the NEAT1 5’ end showed, as expected, abundant NEAT1 foci throughout the nucleus (Figure S1B). We then conducted cell viability assays after sno-NEAT1_1 transfection, and observed a significant increase in the number of sno-NEAT1_1 treated cells compared to the sno-control treated cells (Figure 2E). Overall, these data suggest NEAT1_1 overexpression is associated with increased cell growth in neuroblastoma.

### Development of antisense oligonucleotides to boost formation of paraspeckles in high-risk neuroblastoma cells

Lack of expression of NEAT1_2 in the high-risk, MYCN amplified neuroblastoma cell lines suggests a growth-inhibitory role. To achieve biomimicry of the low-risk, non-MYCN amplified cells, a strategy to increase NEAT1_2 levels and subsequently decrease NEAT1_1 levels in the MYCN amplified cells was devised. We designed several antisense oligonucleotides (ASOs) with the aim of inducing isoform switching from NEAT1_1 to NEAT1_2 (Figure S1C). These ASOs are complementary to motifs at the extreme 3’ end of NEAT1_1, sequences bound by the cleavage and/or polyadenylation machinery needed for producing NEAT1_1 [5]. The rationale was to use the ASOs as steric blockers to interfere with cleavage and/or polyadenylation leading to increased NEAT1_2 at the expense of NEAT1_1. The sequences were designed relative to the PAS, set at ‘0’ in our schematic (Figure S1C). ASOs of 2’O-methyl (2’OMe) chemistry were first used to evaluate effectiveness in U2OS cells (Figure S1D). Two sequences of ASOs (−19 +5, and -14+5) exhibited greatest NEAT1_2 increase, and were selected for further assessment in the neuroblastoma cell lines. Given potential non-specific interactions between paraspeckle proteins and 2’OMe ASOs [45], we switched to morpholino (PMO) chemistry for the remainder of the study. To efficiently transfect PMOs, a complimentary ‘leash’ DNA oligonucleotide was added to allow cationic lipoplex formation [46] (Figure S1E).

High-risk neuroblastoma cells were treated with the two selected ASOs (Figure 3A). FISH against NEAT1_2 in cells treated with each PMO at 25 nM final concentration showed an increase in paraspeckle number in both KELLY (Figure 3B and 3D) and BE2C (Figure S2A) cells compared to control PMO treated cells. These two sequences (−19 +5, and -14+5,) are henceforth called Boost Paraspeckle (BoostPS) 1 and 2 respectively for the remainder of the study (Figure 3A). 25 nM BoostPS 1 PMO increased NEAT1_2 by up to 3.8 fold and 1.8 fold in KELLY and BE2C cell lines respectively (Figure 3C). 25 nM BoostPS 2 PMO increased NEAT1_2 up to 5.1 fold and 2.1 fold in KELLY and BE2C cell lines, respectively (Figure 3C). NONO immunofluorescence, coupled with FISH, showed co-localisation to the paraspeckles, indicating proper paraspeckle formation (Figure 3E and 3F). Therefore, by targeting the PAS site of NEAT1_1 with ASOs, we can increase NEAT1_2 and induce paraspeckles in KELLY and BE2C cells.

### The BoostPS PMOs are capable of forming traditional core-shell paraspeckles, which can sequester paraspeckle proteins

One function of paraspeckles is the subnuclear sequestration of proteins, including NONO, thereby lowering nucleoplasmic levels and inhibiting the downstream target genes of these proteins [27,47,48]. To determine NONO sequestration, we quantified the NONO immunofluorescence signal in the experiments with NEAT1_2 FISH (Fig 3E). We observed an increase in the proportion of NONO within paraspeckles up to 5 fold in the BoostPS PMO treated KELLY cells, with a corresponding depletion of nucleoplasmic NONO (Figure 3F). Thus, BoostPS PMO treatment can increase NONO subnuclear sequestration to paraspeckles in KELLY cells.

Functional paraspeckles have a distinct core-shell ultrastructure, with the 5’-end and 3’-ends of NEAT1_2 in the shell and the middle of NEAT1_2 in the core [16,49]. We conducted structured illumination super-resolution microscopy (SIM) on 50 nM PMO transfected KELLY cells with FISH against the 5’-end of both NEAT1 isoforms (red) and probes targeted to NEAT1_2 middle (green) (Figure 4A). We observed the core-shell structure only in the BoostPS PMO samples, but not in the control (Figure 4B, videos S1, S2 and S3). Thus, KELLY cells do not normally form core-shell paraspeckles, but increasing NEAT1_2 levels with the PMOs can reconstitute this.

### BoostPS ASOs in high-risk cells hinder cell viability and microspeckle formation

We next observed the effects of BoostPS PMOs on the cell growth of the high-risk, MYCN amplified cells. The 25 nM BoostPS 1 and 2 PMO transfected KELLY and BE2C cells had reduced cell confluence and cell viability compared to the control (Figure 5A; further observed in videos S4, S5 and S6). We also tested the PMOs in SK-N-AS (shown above to be unchanged with respect to NEAT1_2 and paraspeckles) and, as expected, observed no difference in cell viability with BoostPS PMO treatment (Figure S2B).

To determine whether the BoostPS PMOs are effective in other cancer contexts, we expanded the use of this treatment to other cancer cell lines. In both A549 lung cancer and HCT116 colorectal cancer cell lines, an increase in NEAT1_2 RNA levels as well as a reduction in cell viability was observed in cells treated with BoostPS 1 PMO compared to the Control PMO (Figure S2C and S2D). These data indicate that isoform switching from NEAT1_1 to NEAT1_2 can occur successfully, accompanied with a reduction in cell viability, in other cancer contexts.

We next examined NEAT1_1 in PMO transfected cells. FISH using the combined red and green probes that target different regions of NEAT1 (as per Figure 4A) in 50 nM BoostPS PMO transfected KELLY cells (Figure 5B) showed a significant reduction in microspeckles (Figure 5C). We confirmed this reduction in NEAT1_1 using RT-qPCR; additionally, for these RT-qPCR experiments extra heating and vortexing steps in the RNA extraction process were undertaken to ensure complete extraction of NEAT1_2, as NEAT1_2 is a semi-extractible lncRNA, and cannot be entirely extracted in traditional RNA isolations [50]. Using comparative quantitation between qPCR amplicons of the NEAT1_2 and total NEAT1_1 products, an increase in NEAT1_2 in BoostPS samples is coupled with subsequent decrease of calculated NEAT1_1 levels (Figure 5D). These combined imaging and qPCR data show that a reduction in NEAT1_1 levels occurs after BoostPS PMO treatment.

To address if the reduction of NEAT1_1 after BoostPS treatment contributed to the reduced cell growth phenotype of treated KELLY cells, we conducted BrdU incorporation assays with KELLY cells transfected with Control or BoostPS 1 PMOs, co-transfected with either sno-control or sno-NEAT1_1 plasmids (Figure 5E, S2E and S2F). A ‘rescue’ of cell viability in BoostPS transfected cells with NEAT1_1 overexpression was observed. These data suggest that the BoostPS PMOs alter cell phenotype in this context, at least in part, by inducing lower NEAT1_1 levels. These data further support the hypothesis that NEAT1_1 has a cell-proliferative role in many cancers, including neuroblastoma.

### BoostPS ASOs upregulate differentiation pathways in MYCN amplified high-risk neuroblastoma cells

In order to assess how modulating NEAT1_1:NEAT1_2 ratios may be affecting multiple cellular pathways, RNA-sequencing (RNA-seq) was conducted on KELLY cells transfected with a control PMO and the BoostPS 1 and 2 PMOs. For differential gene expression analysis, the control oligonucleotide was compared to a combined set of BoostPS 1 and PS 2. In this comparison, 1,313 genes were differentially expressed (Figure 6A). In order to visualise whether the oligonucleotides exerted similar effects, all genes that were differentially expressed in any comparison were plotted as a change from control (Figure 6B). Samples were clustered into oligonucleotide groups, and showed that despite fold change differences, in general, differentially expressed genes changed in the same direction (Figure 6B). To assess whether the observed changes in gene expression were concentrated in specific biological pathways, gene ontology (GO) analysis was performed using Gene Set Enrichment Analysis (GSEA). Significantly enriched pathways primarily resulted from increased gene expression and related to differentiation (Figure 6C). A network consisting of significantly enriched biological processes shows significant functional overlap, which we classified into larger subgroups (Figure 6D). Thus, overall, it appears that BoostPS PMO treatment upregulates differentiation processes.

**Figure 6:**
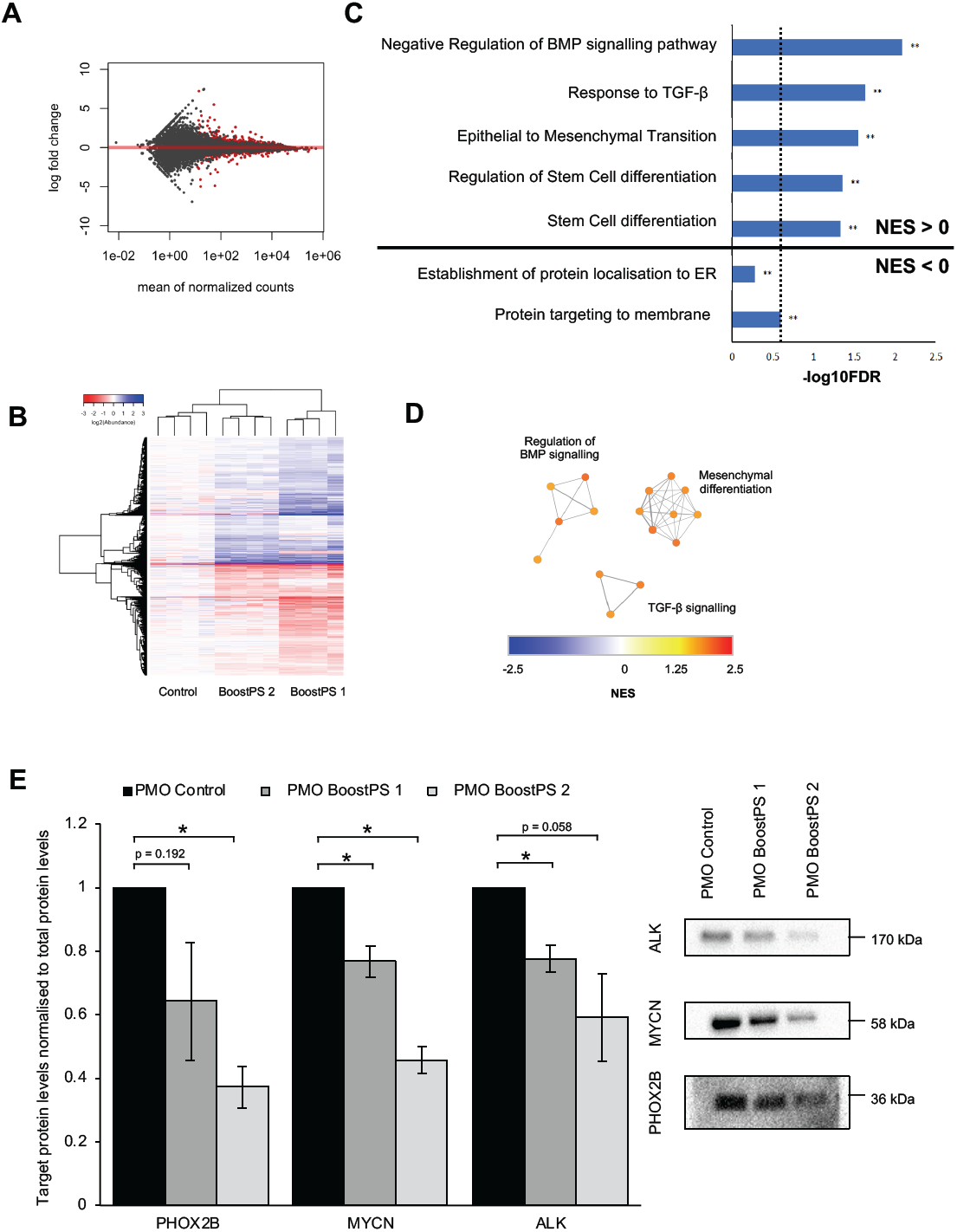
BoostPS PMO treatment results in widespread gene expression changes and upregulation of differentiation processes. (A) Mean gene counts in RNA-seq versus log fold change in a comparison between four control samples (PMO Control) and eight oligonucleotide treated KELLY samples (BoostPS 1 and PS 2). (B) Log2(abundance) calculated as a change from control for all genes that were differentially expressed in any of: control vs BoostPS1, control vs BoostPS2 and control vs BoostPS1 and PS2. (C) Summary of select GO terms from GSEA analysis of control vs BoostPS1 and PS2, using the GO: biological processes gene set. NES = Normalised Enrichment Score, NES > 0 indicates processes that were upregulated with PMO treatment. (D) Network of significant (P,0.001, FDR < 0.1) GO terms coloured by GSEA NES and connected if >35% of genes are shared. Network titles are a summary of observed nodes. (E) Western Blot quantitation analyses of the levels of PHOX2B, MYCN and ALK proteins in KELLY cells treated with 50 nM Control or BoostPS PMOs (n = 3 biological replicates). At right is 1 representative blot (L to R; PMO Control, BoostPS 1 and BoostPS 2. (*P<0.05, **P<0.001, ***P<0.0001)

The induction of differentiation in neuroblastic cells as a therapeutic avenue in neuroblastoma has been thoroughly explored in the past [51,52]. PHOX2B, MYCN and ALK are protein biomarkers indicative of high-risk neuroblastic phenotype and have known roles in maintenance of the pluripotent neuroblastic state in high-risk neuroblastoma cells [37,53,54]. To test if these neuroblastic biomarkers were altered in BoostPS samples, we carried out Western Blotting of treated lysates and observed a decrease in protein expression levels of these three proteins after 50 nM BoostPS PMO transfection in both KELLY (Figure 6E, Figure S3B) and BE2C cells (Figure S3A and S3C). Thus, these data, combined with the RNA-seq data, indicate that modulating the NEAT1_1:NEAT1_2 ratio and increasing paraspeckles is associated with increased differentiation of MYCN amplified, high-risk neuroblastoma cells.

## DISCUSSION

In this study, we have developed ASOs to transiently induce switching of the lncRNA NEAT1_1 isoform to its counterpart, NEAT1_2, increasing the formation of NEAT1_2-dependent paraspeckles. High-risk neuroblastoma cells were targeted due to our finding of their low paraspeckle abundance in high-risk phenotype, and the inverse correlation of NEAT1 with MYCN in neuroblastoma patients. The ASOs reduced NEAT1_1, as well as inducing paraspeckles. Changing NEAT1 ratios in MYCN amplified cells reduced cell proliferation and altered regulation of differentiation pathways, suggesting suppression of NEAT1_2/paraspeckles may maintain pluripotency. To the best of our knowledge, this is the first study involving ASO-mediated lncRNA isoform switching. A further finding with therapeutic implications is that the lncRNAs transcribed from the same gene seem to have opposing roles in the same cancer context.

Numerous studies use typical gapmer ASOs, which induce RNaseH mediated degradation of ASO bound targets [42], to target and downregulate lncRNAs in cancer [55]. However, to our knowledge, our study is the first involving non-gapmer ASOs binding to lncRNAs to alter PAS and RNA processing in cancer. A further example of isoform-switching ASOs with potential as anti-cancer therapeutics, although not involving PAS selection, are those that block the exon 2 splice site of the protein-coding BCL-X gene, resulting in reduced expression of the longer, anti-apoptotic BCL-XL mRNA and increased expression of the shorter, pro-apoptotic BCL-XS mRNA isoform [56]. Although use of RNA therapeutics in cancer is still being developed, in other disease contexts, splicing-modulator ASOs have been FDA-approved for the neurodegenerative conditions spinal muscular atrophy and Duchenne muscular dystrophy [40,41]. Thus, our proof of principle study opens the door to further exploration of polyadenylation-modulating ASOs in both coding and non-coding genes that can be considered as disease therapeutics. More broadly, the potential to modulate lncRNA function with ASOs (as opposed to degrading them) is only starting to be explored. Besides NEAT1, other targets for ASOs could include scaffolding lncRNAs such as HOTAIR, with ASOs designed to block binding of epigenetic modifying proteins, or those lncRNAs that ‘sponge’ tumour-suppressive microRNAs, with ASOs designed to block miRNA binding [57,58].

NEAT1 is well established as one of the most common cancer-associated lncRNAs, although its role as a clear cancer driver is still uncertain. Our observations of NEAT1 as a potential tumour suppressor are at odds with most literature indicating NEAT1 as an oncogene, as seen in meta-analysis and other studies linking higher NEAT1 levels with cancer severity, progression and poor outcome [7]. However, most of these studies do not distinguish between NEAT1 isoforms. Studies using various types of NEAT1 knockout mice, in which one or both NEAT1 isoform have been deleted, have also yielded varying results in cancer models, with one study indicating NEAT1 was required for cellular transformation and tumour growth [22] and another indicating NEAT1 prevented tumour formation [24]. Overexpression of NEAT1_1 partially rescued the tumour-associated phenotype in one model [24], yet NEAT1_1 appears to play no role in a different cancer model [59]. Whilst our analysis in neuroblastoma indicates that high NEAT1 is associated with better outcome, a recent study in a selection of matched cell lines indicates NEAT1 levels can increase in chemo-resistant neuroblastoma tumours, however NEAT1 isoforms were not separated [60]. Hence, future studies into NEAT1 should take care to examine the different roles of the two isoforms and be mindful that NEAT1 roles may vary in different biological and disease contexts. In this regard, the BoostPS ASOs have added to the repertoire of NEAT1 modulating techniques and should be useful for teasing out these roles.

There are likely additional sites in NEAT1 that could also be explored as ASO targets. Additional regions of NEAT1 may be involved in determining the NEAT1_1:NEAT1_2 ratio, as deletion of stretches spanning 2.8 to 4 kb or 5.1 to 5.9 kb increased NEAT1_2 at the expense of NEAT1_1 [49]. Further, deletion of a 100-nucleotide GU-rich TDP-43 binding site only 105 bases upstream of the PAS site also altered the ratio in favour of NEAT1_2 [29]. Of note, future application of the ASOs developed here may be limited to cells in which basal expression of NEAT1_2 is lower than that of NEAT1_1. Although a high NEAT1_1:NEAT1_2 ratio may be the norm in most cells within tissues [61], in cell lines there is evidence of widely varying NEAT1_1:NEAT1_2 ratios [50].

We explored the use of these ASOs in the context of neuroblastoma, as we discovered distinct differences in NEAT1 ratios in different neuroblastoma subtypes. We observed reduced cell viability once the cells displayed increased paraspeckles and reduced NEAT1_1. Given the correlation between high NONO levels and poor outcome in neuroblastoma [31], it is possible that increased sequestration of NONO by the increased paraspeckles shown in this study is partly responsible for the reduction in cell viability, as well as the switch to a more differentiated phenotype, however further experiments are needed to test this possibility. Thus, NONO sequestration, NEAT1_1 reduction, or other possible mechanisms may all explain how paraspeckle increase as a result of ASO treatment may be responsible, at least in part, for regulating transcripts associated with a more differentiated state in the neuroblastoma cells. Future experiments using NONO, NEAT1_1 or total NEAT1 knockout neuroblastoma cells could test this hypothesis.

We do not know the mechanism used by MYCN amplified cells to maintain low transcription of NEAT1 overall, or high NEAT1_1:NEAT1_2 ratio. A recent study showed MYC represses NEAT1 transcription in chronic myeloid leukemia, thus future experiments could test if MYCN acts in a similar fashion in neuroblastoma [62]. Interestingly, distinct stressors can both alter NEAT1 transcription, as well as altering NEAT1_1:NEAT1_2 ratios [18]. For example, mitochondrial stress decreases ATF2, and reduces ATF2-dependent NEAT1 transcription, yet the overall levels of NEAT1_2 increase, suggesting additional post-transcriptional control mechanisms. It is possible that these additional, as yet unknown, mechanisms are suppressed in MYCN amplified, high-risk cells to maintain a high NEAT1_1:NEAT1_2 ratio and suppress paraspeckle production.

We observed that BoostPS ASO treatment led to upregulation of differentiation pathways in MYCN amplified high-risk neuroblastoma cells. This is not altogether surprising, as multiple studies show that there is a correlation between increased paraspeckles and differentiation, and in some settings paraspeckles drive differentiation [29,63,15,4]. The use of differentiation-inducing agents in high-risk neuroblastoma, in which cells remain undifferentiated, is a therapeutic avenue being explored [64], including retinoic acid based differentiation therapy [65]. However, drugs that exploit patient- and tumour-specific alterations that cause the undifferentiated, self-proliferative state are required, in order to develop targeted treatments that would be more likely to succeed. Although our study is a proof of principle, the BoostPS ASOs could be one such potential drug, used in personalised treatment strategies in high-risk neuroblastoma patients.

Although we tested mostly in neuroblastoma cells, the effectiveness of BoostPS ASOs in lung and colorectal cancer cell lines demonstrated that these ASOs may have therapeutic potential in other disease contexts. Perhaps most promising in colorectal cancer, where higher NEAT1_1 expression was correlated with poor patient outcomes and high expression of NEAT1_2 correlated with better patient outcomes [21]. Beyond cancer, BoostPS ASOs could be expanded to other disease contexts such as the neurodegenerative conditions ALS and Huntington’s (HD). NEAT1_2, which is rarely detected in the human central nervous system (CNS), exhibits increased expression in ALS and HD patient CNS samples, with increased paraspeckle formation conferring protective properties [66,67,8].

Overall, the NEAT1 ASOs we have developed can be used as a tool to study the biochemical and functional differences between NEAT1_1 and NEAT1_2, and have therapeutic potential in a variety of diseases. Increased sequence optimisation targeting PAS-independent sites could provide further biochemical information about the NEAT1 transcript, and help dissect the roles of the two NEAT1 isoforms. Future directions may also involve deciphering the role paraspeckles have in altering differentiation pathways in this cancer context. More broadly, this study indicates that steric blocker ASOs to modulate polyadenylation events may be an option to manipulate lncRNAs in cancer and other diseases.

## ACKNOWLEDGEMENTS

We acknowledge the Centre for Microscopy, Characterisation and Analysis at The University of Western Australia for SIM microscopy. We thank other members of the Fox lab for helpful discussions about the manuscript. This work was supported by a research grant from the Cancer Council of Western Australia to AHF and RL [APP1126667] and AHF [APP1106644], a Cancer Council of Western Australia Honours Scholarship (2017) to JALC and a grant from the National Health and Medical Research Council of Australia [APP1147496] to AHF and SF.

## ABBREVIATIONS

NEAT1: Nuclear Paraspeckle Assembly Transcript 1;
NONO: Non-POU domain-containing octamer-binding protein;
MYCN: V-myc myelocytomatosis viral-related oncogene, neuroblastoma derived ;
BoostPS: Boost Paraspeckle;
PMO: morpholino;
2’OMe: 2’O-methyl.

## DECLARATIONS

### Availability of Data and Materials

R2: Genomics Analysis and Visualization Platform is a web based genomics analysis and visualization application, available at (http://r2.amc.nl).

RNA-sequencing data (Figure 6) can be found on GEO with number GSE133562 and reviewer token: wdqduskcfpmtzed

### Authors’ Contributions

AN conducted all wet-lab experiments associated with the neuroblastoma cell lines, designed the experiments, analysed the data and co-wrote the paper. JALC conducted the bioinformatics analyses, analysed the data and co-wrote the Methods section. RL designed the experiments, conducted work done with the U2OS, A549 and HCT116 cell lines, and contributed intellectually to the paper. AH acquired the SIM images. TL provided the neuroblastoma cell lines, and provided intellectual input. SW and SF designed and provided reagents for the 2’O methyl ASO work and provided intellectual input. AHF designed the experiments, analysed the data and co-wrote the paper.

## Figure Legends

**Supplementary Figure 1:**
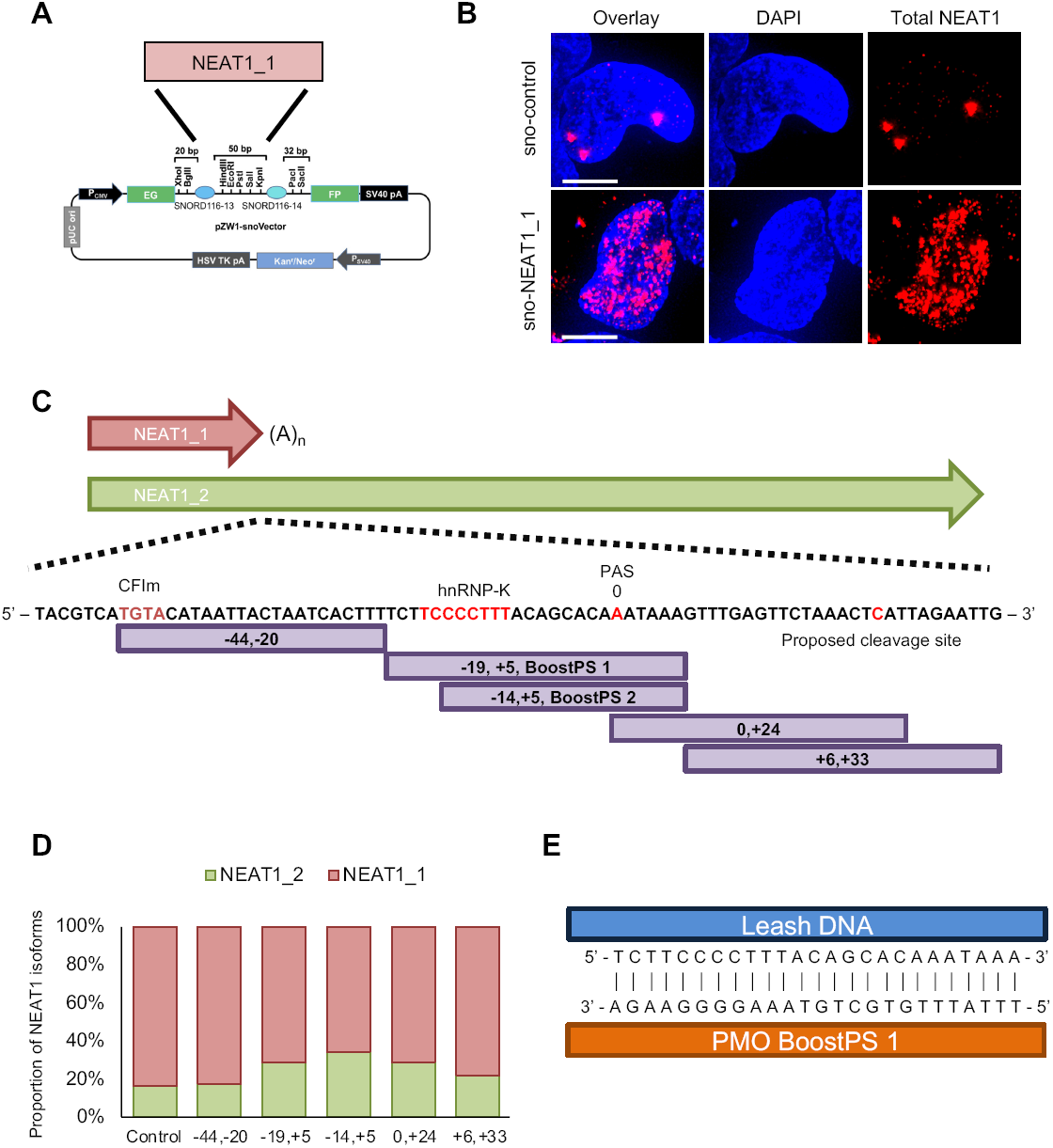
Functional analysis of NEAT1_1 and NEAT1_2 isoforms in high-risk neuroblastoma. (A) Schematic of the sno vector plasmid with human NEAT1_1 in the multi-cloning site. (B) Fluorescence micrograph images of representative cells stained for total NEAT1 (red) in KELLY cells transfected with sno-control plasmid (left), or sno-NEAT1_1 plasmid (right). DAPI (blue) stain indicates cell nuclei. Scale Bar: 5µm. (C) Schematic showing the binding sites for the 2’OMe oligonucleotides, that were designed to cover the region essential for isoform switching. The polyadenylation start site, HNRNP-K and cleavage factor (CFIm) sites are highlighted in red. (D) RT-qPCR data display relative NEAT1 isoform levels in U2OS cells transfected with 100 nM 2’OMe oligonucleotides (n = 1). (E) Schematic depicting DNA oligonucleotide leash binding complimentarily to morpholino to allow lipofection of the complex.

**Supplementary Figure 2:**
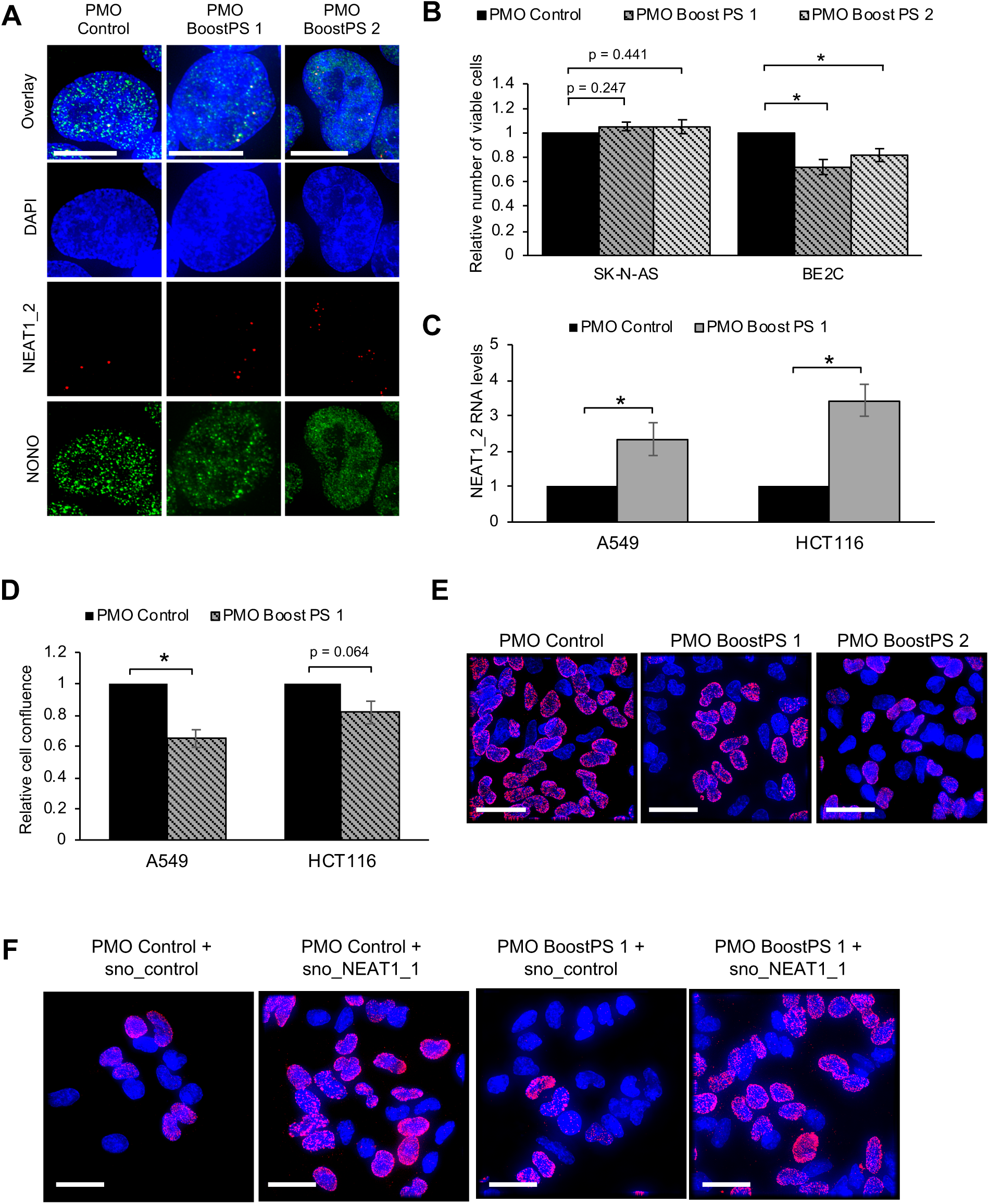
BoostPS PMO-driven cell viability reduction limited only to high-risk neuroblastoma cells, and representative images of BrdU assays. (A) Fluorescence micrograph images of representative cells stained for NEAT1_2 (red) and NONO immunofluorescence (green) in BE2C cells transfected with 25 nM Control or BoostPS PMOs. DAPI (blue) stain indicates cell nuclei. Scale Bar: 15µm. (B) Cell viability was measured using the Cell Titer Glo assay in SK-N-AS and BE2C cells 6 days post-transfection of 25 nM Control or BoostPS PMOs (n = 3). (C) RT-qPCR data indicating relative NEAT1_2 levels in lung and colon cancer cells treated with 25 nM Control or BoostPS 1 PMOs 48 hours post-transfection (n = 3). (D) Relative cell confluence of lung cancer and colon cancer cell lines 3 days post-transfection of 50 nM Boost PS 1 and 2 oligonucleotides (n = 3). (E) Fluorescence micrograph images of representative fields of KELLY cells 4 days after transfection with 50 nM Control or BoostPS PMOs, pulse-labelled with BrdU for 2h, followed by BrdU staining (red). DAPI (blue) stain indicates cell nuclei. Scale Bar: 25µm. (F) Fluorescence micrograph images of representative fields of KELLY cells 4 days after they were co-transfected with 50 nM Control or BoostPS 1 PMO, and either sno-control or sno-NEAT1_1 plasmids pulse-labelled with BrdU for 2h, followed by BrdU staining (red). DAPI (blue) stain indicates cell nuclei. Scale Bar: 25µm. (*P<0.05, **P<0.001, ***P<0.0001).

**Supplementary Figure 3:**
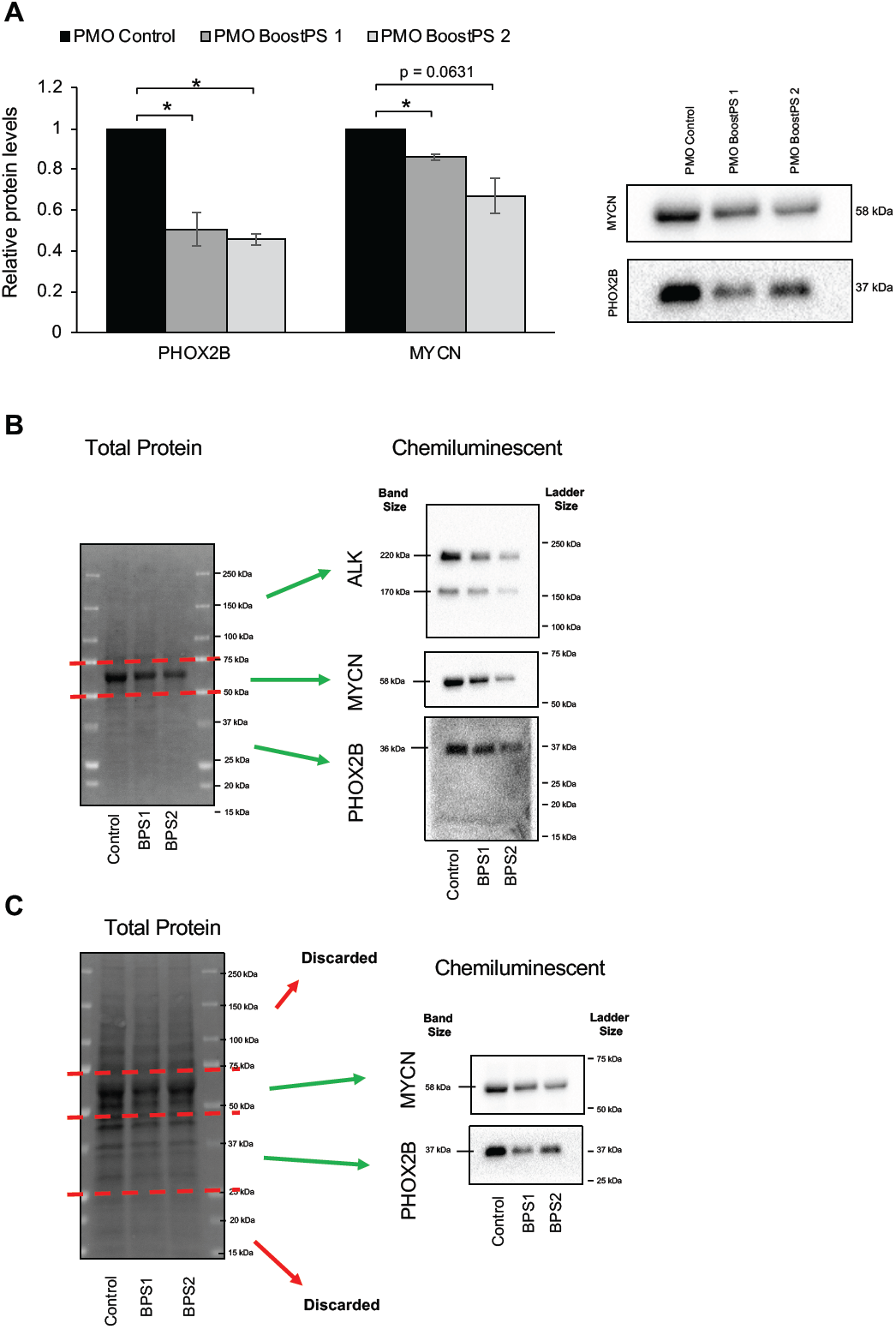
Neuroblastoma protein biomarker levels in BE2C samples treated with BoostPS PMOs, and raw images of Western Blot total protein images and chemiluminescence images. (A) Western Blot analyses show the levels of PHOX2B and N-Myc in BE2C cells 48 h following transfection with 50nM Control or BoostPS PMOs. The blots show 1 representative image, and the graphs depict quantitation of 3 independent experiments or n = 3. (B-C) Uncropped Western Blot data of lysates from (B) KELLY cells, or (C) BE2C cells, transfected with 50nM Control, BoostPS 1, or BoostPS 2 PMOs (left to right) and probed with antibodies against ALK, MYCN, or PHOX2B. In-gel pre-stain allowed for quantification of total protein levels in each lane. Following transfer, the membranes were cut along the red dashed lines. The different membranes were then stained for different antibodies. The total protein values for each lane were used to determine the relative amounts of the target chemiluminescent bands between each of the PMO samples. n = 1. (*P<0.05, **P<0.001, ***P<0.0001)

## Supplementary videos

Video 1 is a 3D rotation of a representative paraspeckle in Control PMO transfected KELLY cells, as marked by FISH against the 5’ end of NEAT1, captured with 3D-SIM.

Video 2 is a 3D rotation of a representative paraspeckle in BoostPS 1 PMO transfected KELLY cells, as marked by FISH against the 5’ end of NEAT1, captured with 3D-SIM.

Video 3 is a 3D rotation of a representative paraspeckle in BoostPS 2 PMO transfected KELLY cells, as marked by FISH against the 5’ end of NEAT1, captured with 3D-SIM.

Video 4 is a longitudinal imaging series over 6 days from the Incucyte capturing KELLY cell confluence following transfection with control PMO.

Video 5 is a longitudinal imaging series over 6 days from the Incucyte capturing KELLY cell confluence following transfection with BoostPS 1 PMO.

Video 6 is a longitudinal imaging series over 6 days from the Incucyte capturing KELLY cell confluence following transfection with BoostPS 2 PMO.

