## Supplemental Table 1 for "PolyA-modulating antisense oligonucleotides reveal opposing functions for long non-coding RNA NEAT1 isoforms in neuroblastoma"

**Supplementary Table 1**

| **Primer Name** | **Sequence** |
| --- | --- |
| RPLP0 F’ | AGC CCA GAA CAC TGG TCT C |
| RPLP0 R’ | ACT CAG GAT TTC AAT GGT GCC |
| Total NEAT1 F’ | GTG GCT GTT GGA GTC GGT AT |
| Total NEAT1 R’ | TAA CAA ACC ACG GTC CAT GA |
| NEAT1_2 F’ | GTC TTT CCA TCC ACT CAC GTC TAT TT |
| NEAT1_2 R’ | GTA CTC TGT GAT GGG GTA GTC AGT CAG |
| MYCN F’ | CGA CCA CAA GGC CCT CAG TA |
| MYCN R’ | CAG CCT TGG TGT TGG AGG AG |
